# Evaluation of longitudinal time-lapsed *in vivo* micro-CT for monitoring fracture healing in mouse femur defect models

**DOI:** 10.1101/692343

**Authors:** Esther Wehrle, Duncan C Tourolle né Betts, Gisela A Kuhn, Ariane C Scheuren, Sandra Hofmann, Ralph Müller

## Abstract

Longitudinal *in vivo* micro-computed tomography (micro-CT) is of interest to non-invasively capture the healing process of individual animals in preclinical fracture healing studies. However, it is not known whether longitudinal imaging itself has an impact on callus formation and remodeling. In this study, a scan group received weekly micro-CT measurements (week 0-6), whereas controls were only scanned post-operatively and at week 5 and 6. Registration of consecutive scans using a branching scheme (bridged vs. unbridged defect) combined with a two-threshold approach enabled assessment of localized bone turnover and mineralization kinetics relevant for monitoring callus remodeling. Weekly micro-CT application did not significantly change any of the assessed callus parameters in the defect and periosteal volumes. This was supported by histomorphometry showing only small amounts of cartilage residuals in both groups, indicating progression towards the end of the healing period. Also, immunohistochemical staining of Sclerostin, previously associated with mediating adverse radiation effects on bone, did not reveal differences between groups.

The established longitudinal *in vivo* micro-CT-based approach allows monitoring of healing phases in mouse femur defect models without significant effects of anesthesia, handling and radiation on callus properties. Therefore, this study supports application of longitudinal *in vivo* micro-CT for healing-phase-specific monitoring of fracture repair in mice.

## Introduction

Adequate monitoring and characterization of the healing process is important in preclinical fracture healing studies. One recent approach to non-invasively capture the formation and remodeling of the osseous fracture callus is the repeated application of *in vivo* micro-computed tomography (micro-CT)^1–3^, allowing for three-dimensional assessment of callus structures over time. A further development is the consecutive registration of time-lapsed *in vivo* callus measurements^4^. In a recent study, we developed a method for registration of time-lapsed *in vivo* scans in a femur defect model in mice using a branching scheme (registration of whole scan for bridged defects; separate registration of the two fragments for unbridged defects)^5^, so that we are now able to also assess dynamic parameters such as bone formation and resorption. In combination with two- and multi-threshold approaches applied in recent studies^5,6^, this allows for better understanding of localized bone turnover and mineralization kinetics during callus formation and remodeling. Whereas many studies focused on understanding the early healing phases (inflammation, repair) with the aim of increasing bone formation and achieving earlier bone union, recent studies also indicate a potential to improve fracture healing outcome via modulation of callus remodeling^7–9^. To address this, time-lapsed *in vivo* micro-CT-based monitoring approaches seem particularly suitable, as not only bone formation but also bone resorption can be reliably monitored over time. Furthermore, as each animal can be followed individually throughout the healing process with low variance in the assessed parameters, animal numbers can be reduced compared to well-established cross-sectional studies with endpoint three-dimensional micro-CT and two-dimensional histological callus evaluation.

In non-fractured bone, longitudinal time-lapsed *in vivo* micro-CT has been increasingly used to monitor changes in bone properties associated with different diseases and external factors, e.g. estrogen-deficiency^10–12^, mechanical-(un)loading^13–15^, and drug application^16^. However, several studies indicate that anesthesia, cumulative radiation dosage and stress due to the required handling for the CT measurements may have effects on animal well-being and on trabecular and cortical bone properties^11,17–21^. Isoflurane is the most commonly used inhalation anesthetic in rodents^22^, characterized by a quick on- and offset of anesthesia and low metabolism rate^22^. According to the EU Directive 2010/63, the severity of repeated isoflurane anesthesia can be categorized as mild, although repeated anesthesia was considered worse than a single session with sex-dependent differences in perceiving the severity of a procedure^17^. Specifically, female mice were shown to be more susceptible to anesthesia-induced effects on well-being compared to male mice^17^. Nevertheless, in both sexes repeated isoflurane anesthesia caused only short-term mild distress and impairment of well-being, mainly in the immediate post-anesthetic period^17^. Radiation has also been shown to have dosage-dependent effects on bone cells *in vivo* and *in vitro*: Whereas high dose x-ray radiation (2.5-8Gy) was associated with reduced osteoblast and osteoclast proliferation^23–25^, lower doses (<2Gy) had a stimulatory effect on osteoclasts. Some studies reported radiation-associated effects on structural bone parameters with stronger and earlier effects seen in trabecular bone compared to cortical bone, whereas other studies did not see any changes^11,19,21,26,27^. Similarly, pre- and post-operative radiation treatments have been associated with changes in fracture healing. Whereas high dose x-ray radiation (5-8Gy) was associated with an impairment of fracture healing, lower doses (0.5-1Gy) accelerated endochondral and intramembranous ossification during the repair process^28–31^. These findings indicate the importance of study-specific adaptation of micro-CT protocols, and to protect animals scanning times and radiation settings should be minimized.

Recently, several fracture healing studies have applied longitudinal *in vivo* micro-CT to monitor callus formation under different conditions (e.g. osteoporosis, polytrauma) and treatments (e.g. mechanical loading, pharmaceuticals)^2–4,32,33^. However, none of these studies assessed, whether longitudinal imaging itself has an impact on callus formation and remodeling. As several studies have reported adverse effects of longitudinal imaging during normal bone remodeling (e.g. reduction in trabecular and cortical thickness)^11,19,21,27^, there is a need to also assess imaging-associated impact on the callus formation during the highly metabolically active process of fracture healing.

Therefore, the objectives of this study were to establish an *in vivo* micro-CT based approach for longitudinal monitoring of fracture healing in a mouse femur defect model and to assess the combined effect of radiation, anesthesia and handling associated with weekly time-lapsed micro-CT measurements on callus properties during the remodeling phase of fracture healing.

## Results

In order to enable longitudinal monitoring of fracture healing in single animals, we established a time-lapsed *in vivo* micro-CT based approach for mouse femur defect models. By registration of consecutive scans, structural and dynamic callus parameters can be followed in three callus sub-volumes (defect center, defect periphery, cortical fragment periphery) and the adjacent cortical fragments over time (Figure 1). To capture potential effects of consecutive micro-CT measurements on callus properties, bone parameters of a scan group subjected to weekly measurements between weeks 0 and 6 were compared to control animals, that were scanned only post-operatively (d0) and after 5 and 6 weeks, respectively (for detailed study design see Supplementary Fig. S1 online).

**Figure 1.**
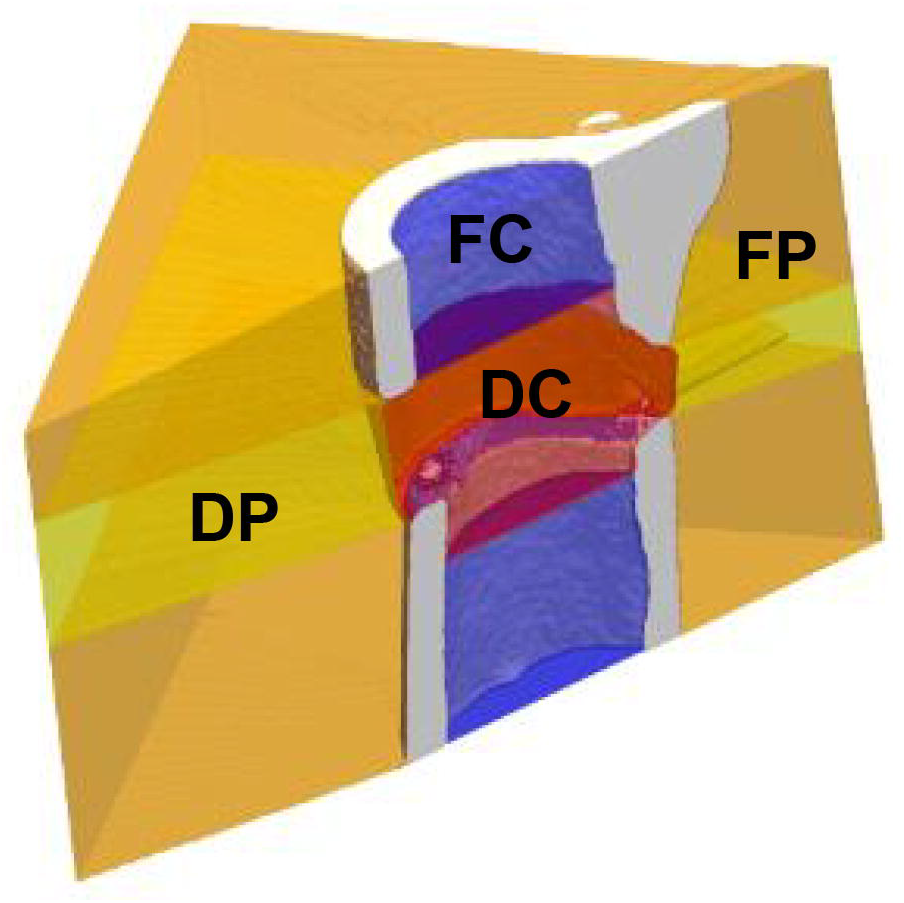
Volumes of interest (VOIs) for micro-CT evaluation of callus and adjacent bone: Defect center (DC - red), defect periphery (DP - yellow), fragment center (FC - blue), fragment periphery (FP - orange).

### General physical observation

All mice recovered rapidly from surgery. During the healing period, in both groups, the animals’ body weight did not significantly change compared to pre-operative values without significant differences between the two groups (see Supplementary Fig. S1 online). We also assessed the social interaction between mice (animals sitting in groups vs. separated from others) and the nesting behavior (start of nest building after surgery). Specifically, all animals started to build nests on the day of surgery and were found in groups in the nest in the morning of post-operative day 1. Throughout the further healing period social interaction between mice and nesting behavior did not differ from pre-surgical observations and was similar for animals of the scan and control groups.

### Volumes of interest (VOI) for evaluation by time-lapsed *in vivo* micro-CT

In order to exclude bias in the further micro-CT analyses, we compared the size of the different VOIs (depicted in Fig. 1) in the control and the scan group. One animal from the scan group could not be included in the analysis due to failure in VOI generation caused by too little cortical bone being present in the field of view (for details on VOI generation see ^5^). The VOIs encompassed the following volume for the control (n=8) and scan group (n=10): 2.51±0.34mm^3^ vs. 2.60±0.34mm^3^ for the defect center (DC), 17.26±2.21mm^3^ vs. 18.73±4.64mm^3^ for the defect periphery (DP), 1.86±0.48mm^3^ vs. 1.73±0.45mm^3^ for the cortical fragments (FC), 14.47±3.30mm^3^ vs. 13.10±2.66mm^3^ for the fragment periphery (FP). The total volume (TOT=DC+DP+FC+FP) between the inner pins of the fixator was 36.10±2.81mm^3^ for the control and 36.15±3.74mm^3^ for the scan group. No significant differences in volume were detected in any of the VOIs between groups.

### Longitudinal monitoring of fracture healing by time-lapsed *in vivo* micro-CT

In the scan group (n=10), the repeated micro-CT scans (1x/week, Fig. 2) covered the period from the day of the defect surgery (d0) until post-operative week 6 with distinct callus characteristics indicative of the different healing phases (inflammation, repair, remodeling; significant weekly differences in bone parameters indicated in Fig. 3). From week 0-1 to week 1-2 a significant 5.7x increase in bone formation was detected (Fig. 3a) in the total VOI (TOT=DC+DF+FC+FP, p<0.0001; Fig. 1), indicating progression from the inflammation to the reparative phase. This led to a significant gain in bone volume by week 2 (BV/TV_week2_: 39±7% vs. BV/TV_week0_: 25±3%, p=0.0134; Fig. 3b). From week 1-2 to week 2-3 a significant 2.8x increase in resorptive activities (p=0.0020; Fig. 3a) was seen, indicating the progression from the repair to the remodeling phase. Two weeks after surgery the highly mineralized bone fraction in the TOT VOI was significantly lower compared to the post-operative measurement (BV_645_/BV_395_ in week 2: 59±5% vs. BV_645_/BV_395_ in week 0: 84±1%, *p*<0.0001), indicating formation of mineralized callus of low density. From post-operative week 2 onwards, the highly mineralized bone fraction in the TOT VOI gradually increased in all subsequent weeks of the healing period reaching statistical significance by week 5 (BV_645_/BV_395_ in week 2: 59±5% vs. BV_645_/BV_395_ in week 5: 79±3%, p=0.0134; Fig. 3c).

**Figure 2.**
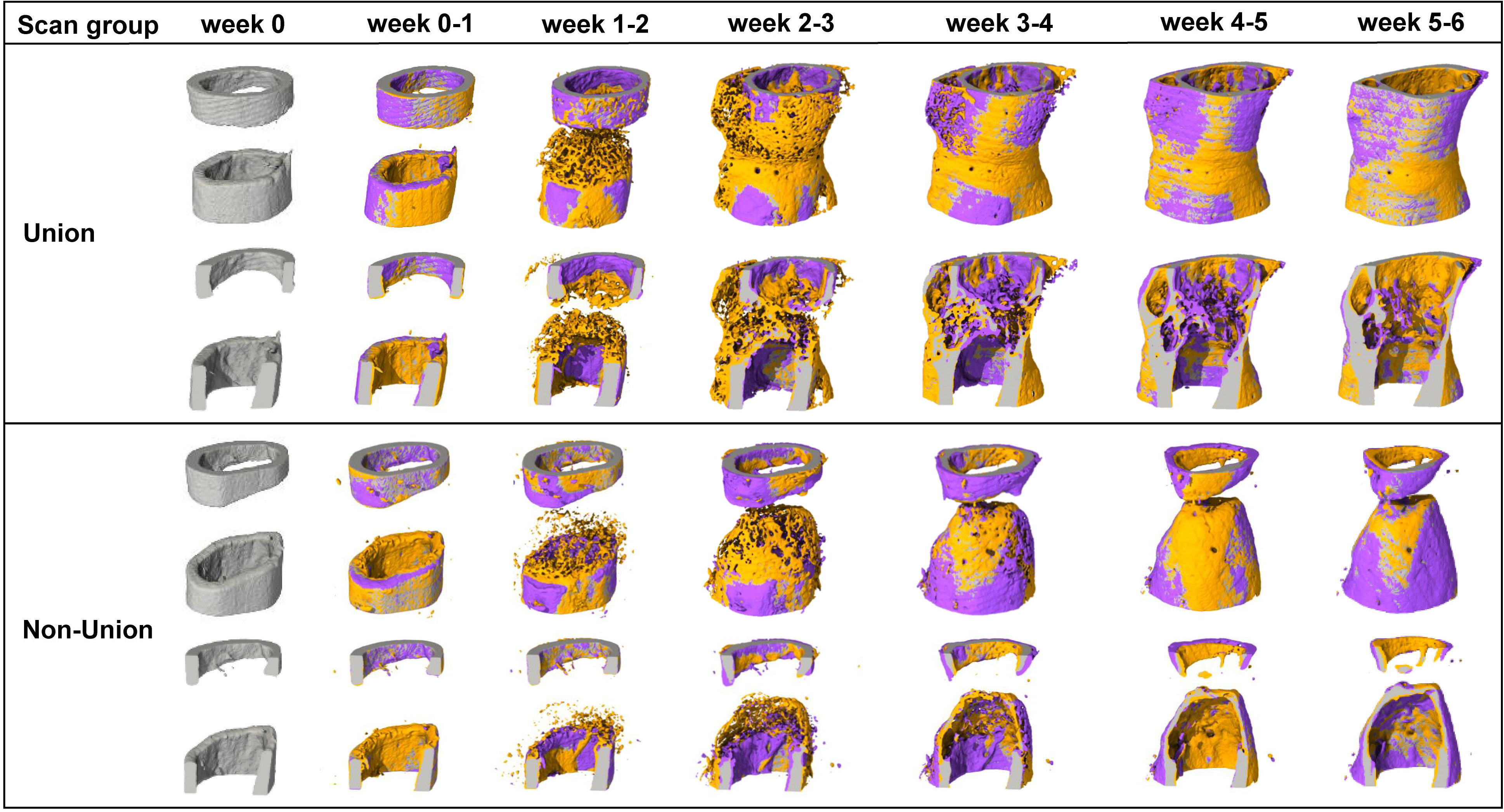
Representative images (full image and cut; threshold: 645 mg HA/cm^3^) of the defect region from animals of the scan group (week 0-6): union defect (top panel), non-union defect (bottom panel). Visualization of bone formation (orange) and resorption (blue) via registration of micro-CT scans from week 5 and 6.

**Figure 3.**
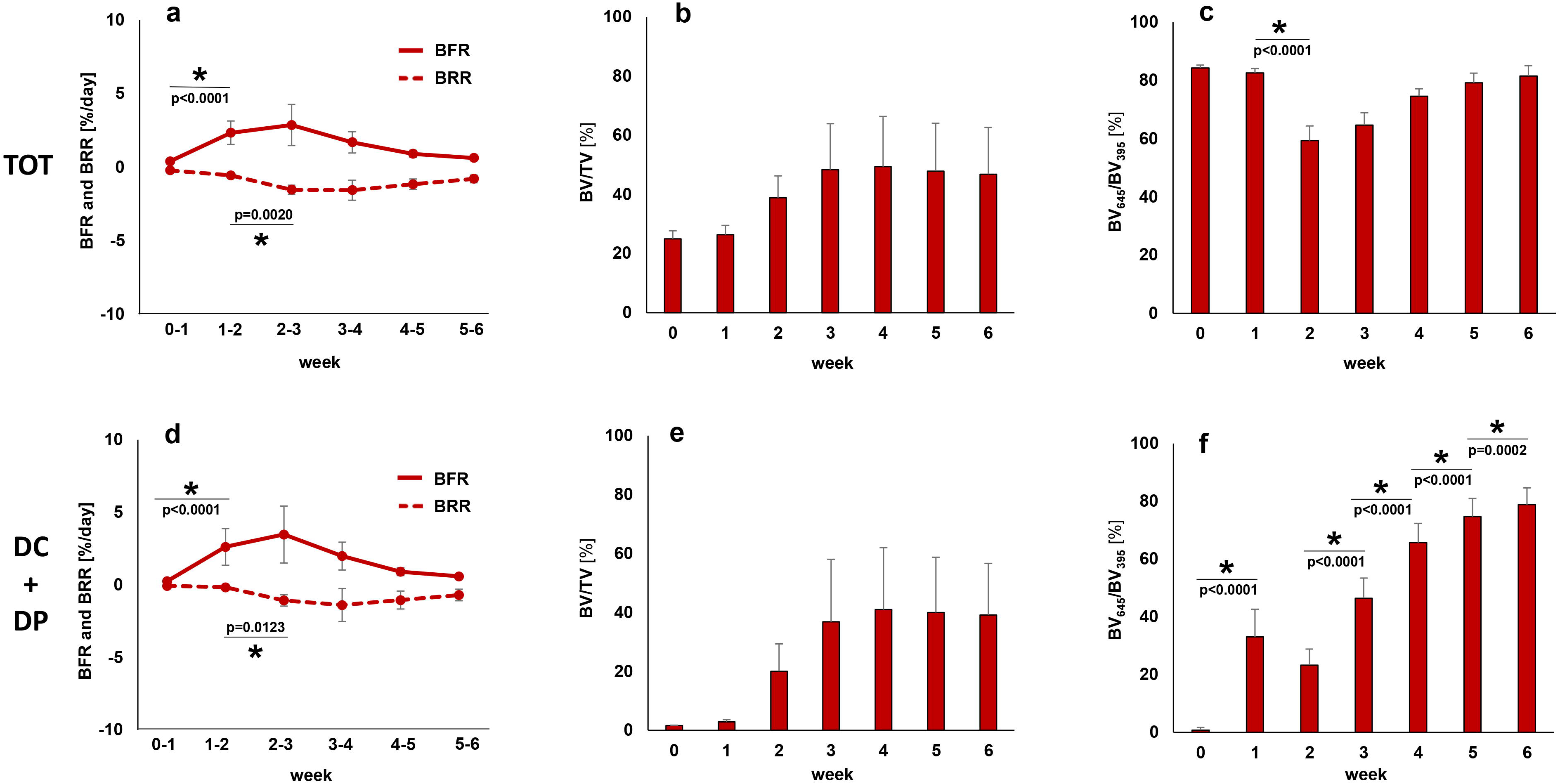
Micro-CT based evaluation of bone parameters in the scan group using different VOIS: total VOI (TOT) - defect and adjacent bone fragments (top), defect VOI (DC+DP) – defect center and periphery (bottom). a+d. Bone formation rate (solid) and bone resorption rate (dashed line) in the femur defect (TV) given in percent per day. b+e: Bone volume (BV) normalized to TV (DC+FC for TOT and DC for DC+DP). c+f: Degree of bone mineralization given as ratio of bone volume with a density ≥645 mg HA/cm^3^ to the total osseous volume in the defect (threshold ≥395 mg HA/cm^3^). n=10; * indicates *p* < 0.05 between consecutive weeks determined by Friedman test with Dunn correction for multiple comparisons (a-e)/ repeated measurements ANOVA with Bonferroni correction (f).

In order to better capture the regions where bone is mainly formed and resorbed, we evaluated the different VOIs separately (Table 1): In the early post-operative phase from week 0-1 to week 1-2 a strong onset of bone formation was seen in the DC and FP sub-volumes, leading to significant 11.8x and 3.4x gain in mineralized tissue from week 0 to week 2 for DC (p=0.0090) and FP (p=0.0091), respectively. This indicates that both intra-cortical as well as periosteal callus formation takes place in this femur defect model. In both VOIs bone formation triggered the initiation of bone resorption from week 2-3. In detail, there was a significant 6.2x and 2.9x increase in bone resorption from week 1-2 to week 2-3 in the DC (p=0.0040) and the FP VOI (p=0.0079), respectively. Compared to the DC and FP VOIs, the initiation of bone formation was much less pronounced in the DP sub-volume with less deposition of mineralized tissue leading to only little peripheral callus formation in this VOI (week 2-6) and subsequently low bone resorption activities from week 2-3 to week 5-6. In all three regions (DC, DP, FP) the fraction of highly mineralized bone considerably increased from week 2 to week 5 (+192%, +774%, +227%), indicating callus maturation. In this femur defect model, callus formation and remodeling mainly took place in the defect region (DC) with only little peripheral callus formation and remodeling (DP). Looking at the cortical fragments (FC), a significant 3.2x increase in resorptive activities was detected in week 1-2 compared to week 0-1 (p=0.0421), whereas no significant weekly change in bone formation activities was seen in this region throughout the assessed healing period. This resulted in a significant 24% reduction in bone volume from week 0 to week 4 (p=0.0091). The fraction of highly mineralized bone also gradually decreased in the remaining osseous tissue reaching statistical significance by week 3 (BV_645_/BV_395_ in week 3 reduced by 9% compared to week 0, p=0.0027).

**Table 1.**
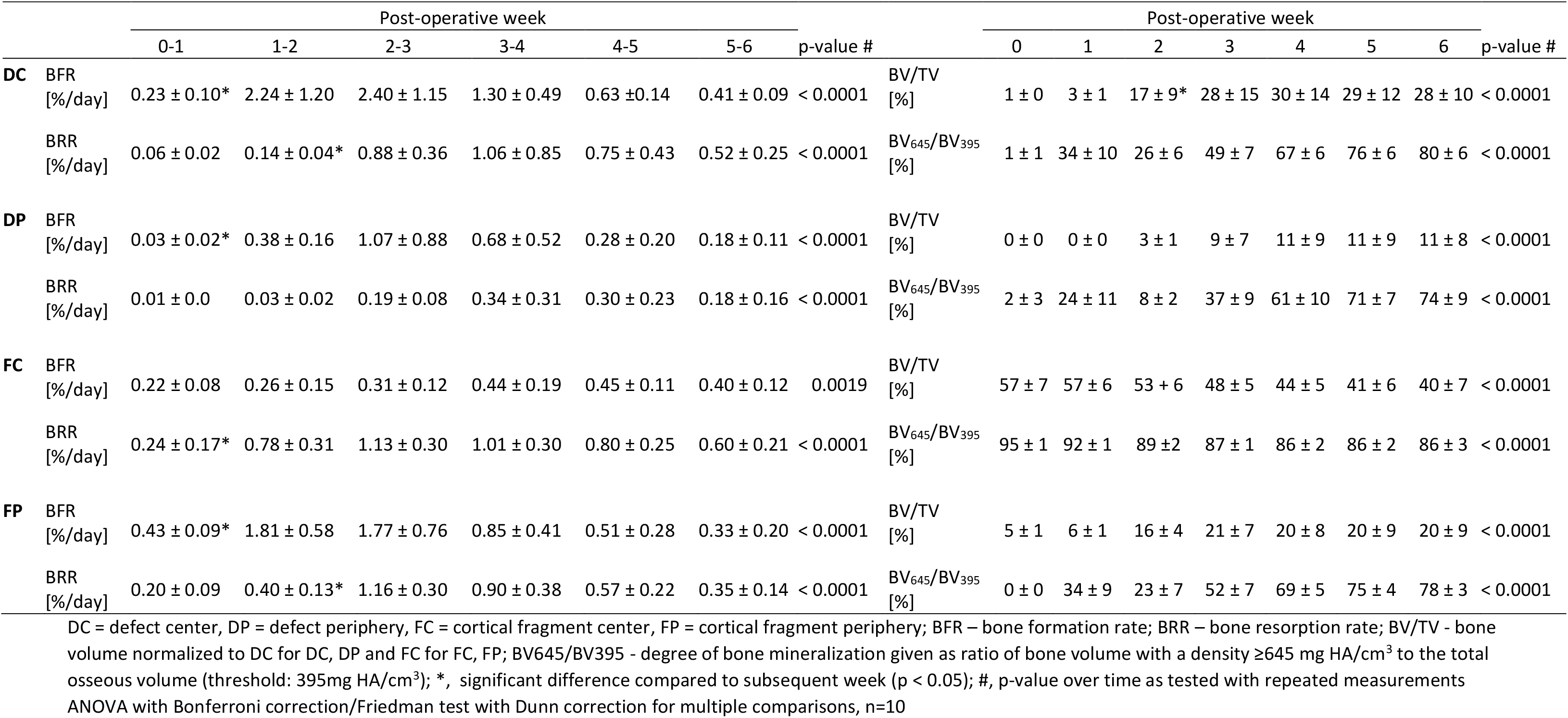
Micro-CT based evaluation of bone parameters in the scan group using different VOIS

### Influence of the longitudinal *in vivo* micro-CT protocol on callus properties

To assess the combined effects of radiation, anesthesia and handling associated with weekly micro-CT measurements (week 0 - week 6) on callus properties, callus parameters of the scan group were compared to control animals that were scanned only directly post-operatively (d0) and after 5 and 6 weeks (Fig. 4). To exclude bias in the further micro-CT analyses, we assessed the post-operative bone volume (BV/TV on d0) and did not see any significant differences between groups: 1.82±0.39% (control) vs. 1.47±0.15% (scan) in DC, 0.16±0.12% (control) vs. 0.13±0.06% (scan) in DP, 56.82±2.31% (control) vs. 57.14±6.99% (scan) in FC, 4.66±0.30% (control) vs. 4.74±0.76% (scan) in FP, 26.92±4.69% (control) and 24.90±2.88% (scan) in TOT.

**Figure 4.**
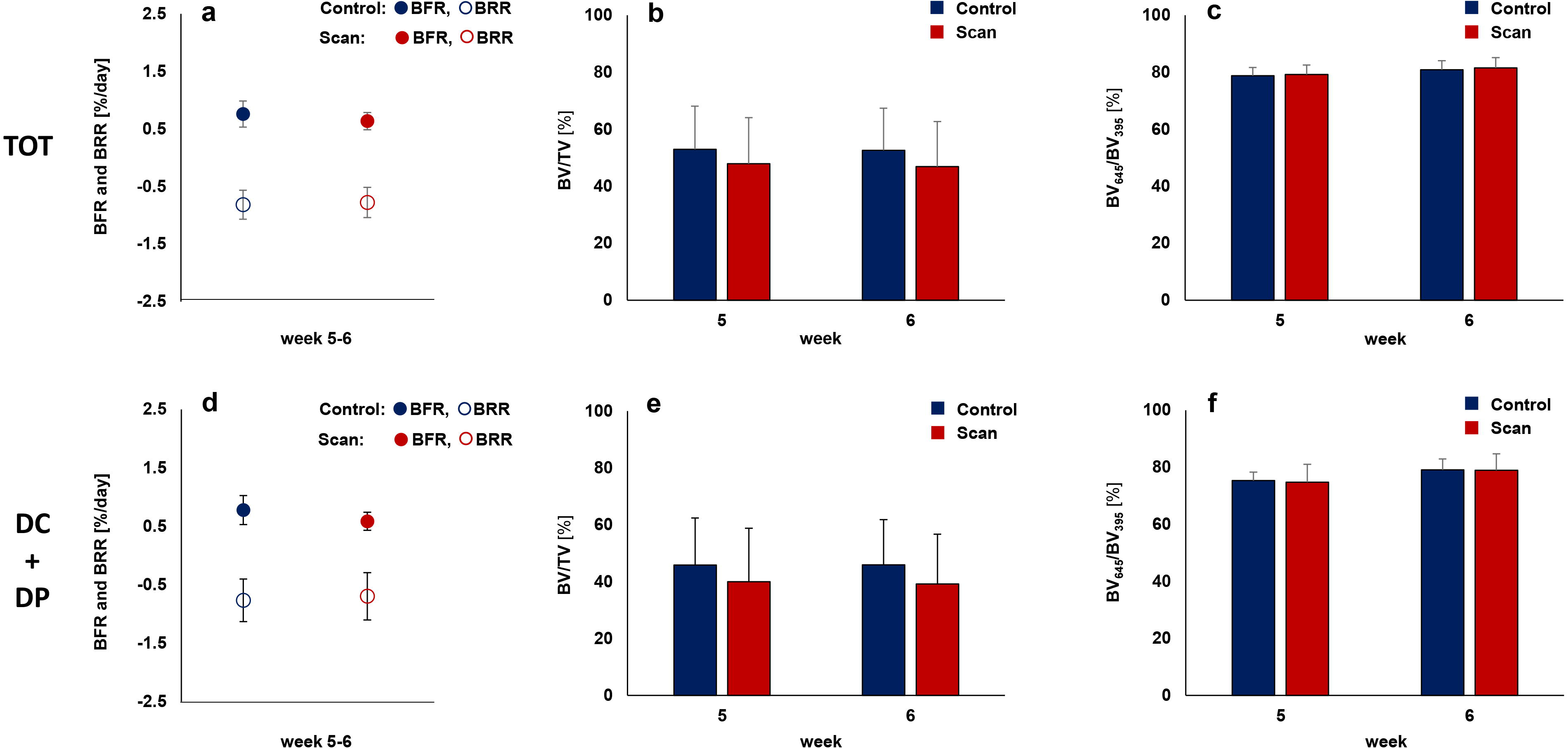
Micro-CT based evaluation of bone parameters in the scan (red) and control group (blue) using different VOIS: total VOI (TOT: DC+DP+FC+FP) - defect and adjacent bone fragments (top), defect VOI (DC+DP) – defect center and periphery (bottom). a+d. Formed (solid) and resorbed (empty) bone volume (BV) in the femur defect (TV) given in percent per day. b+e: Bone volume (BV) normalized to TV (DC+FC for TOT and DC for DC+DP). c+f: Degree of bone mineralization given as ratio of bone volume with a density ≥645 mg HA/cm^3^ to the total osseous volume in the defect (threshold ≥395 mg HA/cm^3^). Control group: n=8, scan group: n=10. * indicates *p* < 0.05 determined by two-tailed Student’s t-test (a-e; f: week 6)/Mann-Whitney U-test (f: week 5).

During the remodeling phase (post-operative weeks 5-6), no significant difference in bone turnover was seen in the total volume (TOT) between the inner pins of the fixator (BFR: 0.76±0.23% in controls, 0.64±0.15% in scan group; BRR: 0.82±0.25% in controls, 0.78±0.26% in scan group). Also, bone volume did not significantly change due to the applied micro-CT protocol with a similar fraction of highly mineralized bone after 5 weeks (BV_645_/BV_395_: 79±3% in controls, 79±3% in scan group) and 6 weeks (BV_645_/BV_395_: 81±3% in controls, 82±4% in scan group).

As radiation effects are known to vary for different bone cell types and locations^30,34,35^, we assessed the four sub-volumes separately (Table 2). The applied micro-CT protocol did not significantly affect bone formation and resorption activities in the callus VOIs (DC, DP and FP) from week 5-6 with similar bone volume observed in week 5 and 6 for controls and scanned animals. In both groups, bone volume remained stable from week 5 to week 6, whereas the density of the mineralized tissue increased during the same period (DC, DP, FP). Looking at the adjacent cortical fragments (FC) also no significant differences were detected in any of the assessed parameters between the control group and the scan group. Similarly to the callus VOIs, bone volume also remained stable in this region from week 5 to week 6. However, in contrast to the callus VOIs, the density of the mineralized tissue in the cortical VOI FC did not change from week 5 to week 6. In week 6, the FC VOI comprised 33% and 35% of the osseous tissue in the total VOI (TOT) for the control and scan group, respectively. In the DC VOI, 37% (control group) and 36% (scan group) of the total bone volume were seen. Less osseous tissue was detected in the two peripheral VOIs, 11% (control group) and 12% (scan group) in the DP and 19% (control group) and 17% (scan group) in the FP VOI.

**Table 2.**
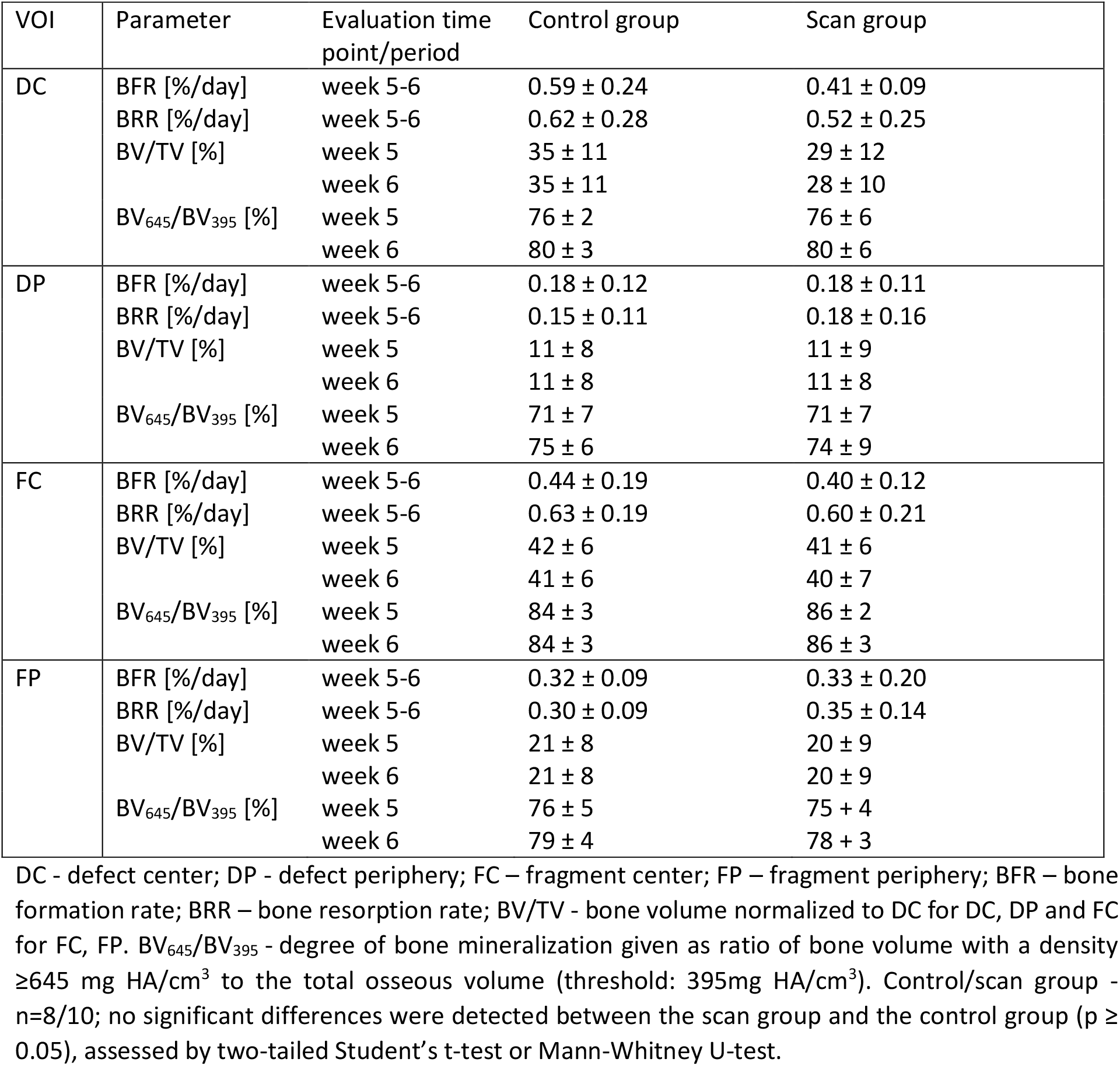
CT evaluation of bone parameters in the different volumes of interest (VOI) for the scan and control group

As group comparisons were only performed from week 5-6, we particularly focused on the defect VOIs (DC+DP) which are most important for evaluating later healing time points during the remodeling phase of fracture healing. No significant differences in bone turnover, bone volume and mineralization were seen between the scan and control group (Figure 4 d-f, Table 2). In addition, according to the standard clinical evaluation of X-rays, the number of bridged cortices per callus was evaluated in two perpendicular planes and animals with ≥3 bridged cortices were categorized as healed. By week 5, cortical bridging occurred in 75% of the control animals and 64% of the scanned animals (Table 3). However, we noticed that non-unions only occurred in defects of ≥1.5mm length. Although the mean defect length (1.45mm±0.16 in controls, 1.47mm±0.16 in scan group) did not significantly differ between groups (p=0.816), the percentage of animals with defects ≥1.5mm was different in the control and the scan group (55% vs. 38%; Table 3). Taking into account only these defects (≥1.5mm length), both groups had a non-union rate of 67% indicating no significant effect of the chosen micro-CT protocol on clinical fracture healing outcome.

**Table 3.**
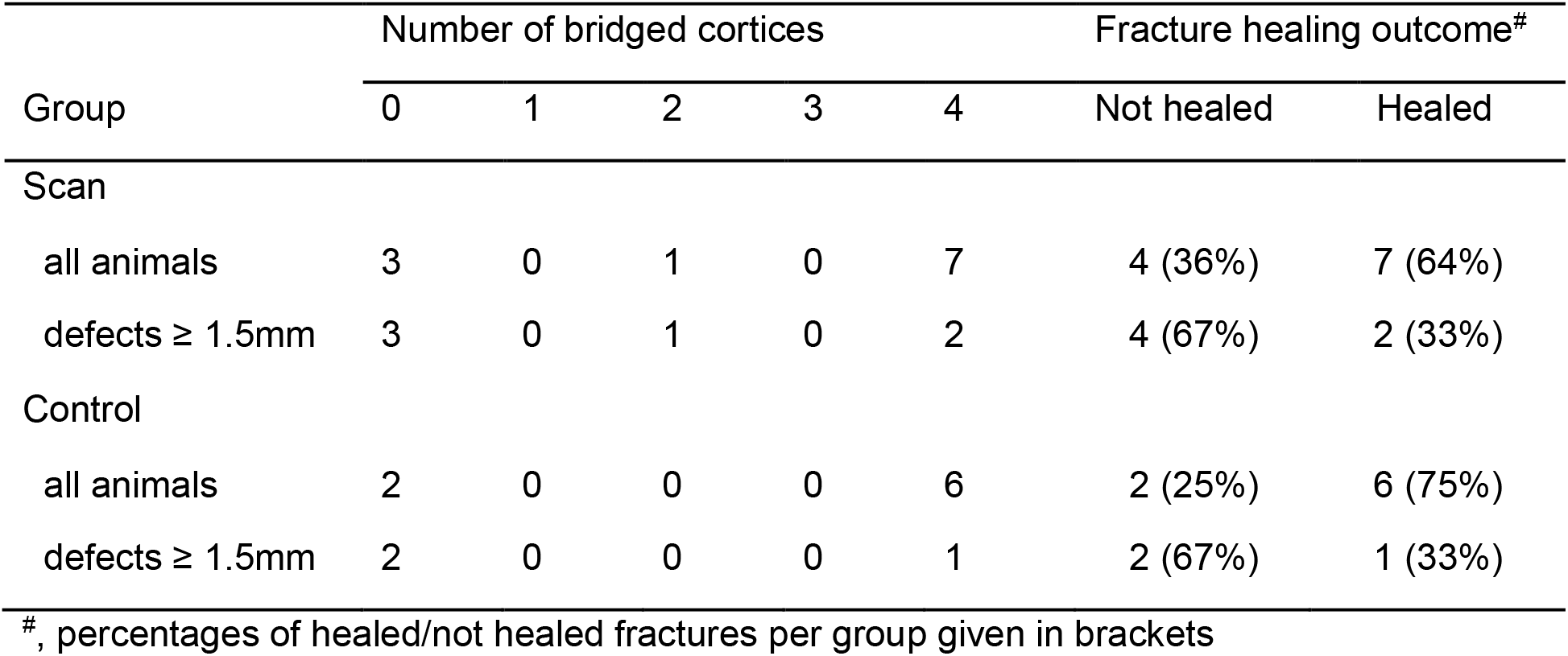
Number of bridged cortices per callus evaluated in two perpendicular planes and number of mice with successful fracture healing (≥ 3 bridged cortices) evaluated 5 and 6 weeks after the defect surgery.

| Group | Number of bridged cortices |  |  |  |  | Fracture healing outcome<sup>#</sup> |  |
| --- | --- | --- | --- | --- | --- | --- | --- |
|  | 0 | 1 | 2 | 3 | 4 | Not healed | Healed |
| Scan |  |  |  |  |  |  |  |
| all animals | 3 | 0 | 1 | 0 | 7 | 4 (36%) | 7 (64%) |
| defects $\geq 1.5$mm | 3 | 0 | 1 | 0 | 2 | 4 (67%) | 2 (33%) |
| Control |  |  |  |  |  |  |  |
| all animals | 2 | 0 | 0 | 0 | 6 | 2 (25%) | 6 (75%) |
| defects $\geq 1.5$mm | 2 | 0 | 0 | 0 | 1 | 2 (67%) | 1 (33%) |

<sup>#</sup>, percentages of healed/not healed fractures per group given in brackets

### Histology

Histomorphometric analysis of Safranin-O stained sections did not show any significant differences in tissue composition (bone, cartilage, fibrous tissue, bone marrow) in the former defect region between the control and the scan group six weeks after osteotomy (Fig. 5, Supplementary Fig. S2 online). The bone tissue fraction was 38+7% and 32+6% for the control and the scan group, respectively. Hardly any cartilage residuals (< 1% of total area) were present in both groups indicating progression of the healing process in the final remodeling stage (Supplementary Fig. S3 online).

**Figure 5.**
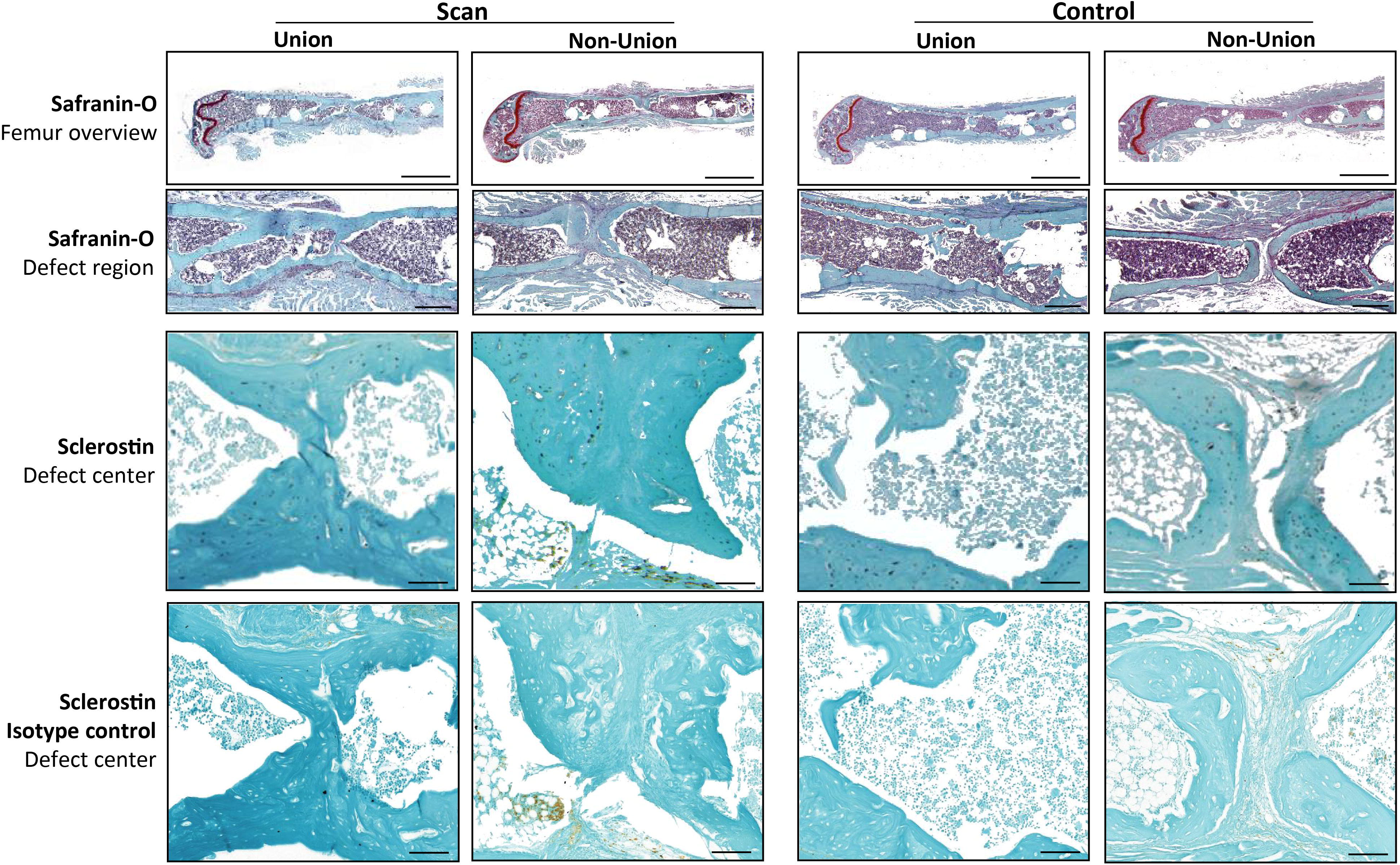
Representative longitudinal sections of fractured femora 6 weeks after defect surgery of unions and non-unions in the scan and control group. Top panel: Safranin-O staining - overview images (row 1), scale bar = 2 mm; area between inner pins of fixator (row 2), scale bar = 500 µm; bottom panel: Sclerostin staining – area centered between inner pins of fixator (row 3), scale bar = 100 µm; Isotype control for Sclerostin staining (row 4), scale bar = 100 µm.

As radiation-mediated decreased bone formation by osteoblasts has previously been associated with the inhibition of osteoanabolic Wnt-signaling^36^, we assessed Sclerostin expression in the defect region by immunohistochemistry. Visual inspection did not reveal any differences in staining intensity and amount of Sclerostin between the scan and control group (Fig. 5).

## Discussion

In this study, longitudinal *in vivo* micro-CT was applied for monitoring the process of fracture healing in mice. In addition, the combined effect of radiation, anesthesia and handling associated with the established imaging approach on callus parameters was assessed. Until now, preclinical fracture healing studies predominantly have a cross-sectional study design with end-point callus evaluation in sub-groups of animals at different time points during the healing period. Only recently, several studies in murine diaphyseal femur defect models stabilized by external fixation^2–4,32^ or intramedullary nails^33^, have applied longitudinal *in vivo* micro-CT to monitor the healing progression in each animal over time. These studies were able to consecutively assess callus parameters (e.g. callus volume and density) at specific time points during the healing process for up to 12 weeks^2–4,32,33^. By registering consecutive scans, we were now able to also include dynamic parameters such as bone formation and resorption in our micro-CT based monitoring approach for fracture healing (for details on method see ^5^). This allows for characterization of the different healing phases seen by changes in formation and resorption in the osseous callus volume. Compared to histology-based determination of the bone formation rate via *in vivo* application of fluorescent dyes (e.g. calcein, alizarin), our micro-CT-based analysis with registration of consecutive scans also allowed for assessment of the bone resorption rate in individual animals. Furthermore, implementation of a two-threshold approach allowed to monitor mineralization kinetics of the fracture callus. Our micro-CT-based approach is therefore particularly suited to better understand callus remodeling under different conditions (e.g. premedication with bisphosponates and associated inhibition of callus remodeling) and to assess pharmacological and mechanical therapies targeting callus remodeling.

Specifically, we saw that the initiation of bone formation (maximum in week 2-3), indicating the onset of the reparative phase, triggered bone resorption (maximum in week 3-4) with maximum osseous callus volumes in week 3. These observations are in line with cross-sectional histomorphometric analyses in similar defect models, where maximum callus area was seen in weeks 2 and 3^37–39^. From week 4 to week 6, bone formation and resorption continuously decreased to levels (BFR ≤ 1%/day; BRR ≤ 0.8%/day) seen in normal bone remodeling of trabecular and cortical bone in mice^14,40–43^, indicating proceeding towards the end of the healing process. Whereas these remodeling activities also led to a decrease in callus volume from week 4 to week 6, the fraction of highly mineralized bone increased during the same period indicating advanced callus maturation. These observations were further supported by histology with no remaining cartilage in the defect region of unions and only a small amount of cartilage residuals in some non-unions. The findings are also in accordance with previous longitudinal studies of up to 4 weeks in mice^2,3,32^ and 8 weeks in rats^4^ mainly focusing on the inflammation and reparative phase. Similar to our study, the bone volume continuously increased during the 4/8 week healing period, although no remodeling to the original bone geometry was seen due to the shorter observation period. One study in ovariectomized rats with a 12-week post-operative monitoring period detected maximum osseous callus volumes by week 6, which diminished thereafter, whereas bone mineral density continuously increased until week 12, similar to our findings. This indicates that during the reparative phase first low-density bone is formed, which is then further mineralized during the remodeling phase of fracture healing. Looking at the functional fracture healing outcome, complete cortical bridging was seen in 64% of the animals, which is similar to other studies (60-62%) using similar femur defect models with relatively stiff external fixation (fixator stiffness measured in our study 24N/mm) in mice^2,44^. These commercially available rigid external fixators are widely used in fracture healing studies as they also allow for the assessment of biomaterials in non-union defects (induced by increasing the defect size). Borgiani et al.^37^ have recently extensively characterized the healing process (week 1-3) using external fixators of similar stiffness and observed endochondral ossification processes, albeit less pronounced compared to semi-rigid fixation. When comparing the outcome of these studies using rigid fixation, it is important to consider differences in defect sizes: 0.7mm^2^, 1.19±0.25 mm^44^, 1.47±0.16mm (this study). In contrast to the other studies, we saw the manifestation of non-unions only in defects ≥1.5mm. This might be partially due to the fact that in some studies the exact defect length might have differed from the reported value (saw diameter used for osteotomy) and that non-union formation was assessed irrespective of defect length. Overall, all longitudinal fracture healing studies using *in vivo* micro-CT were able to follow the healing progression in single animals, thereby reducing variance of outcome parameters compared to cross-sectional studies, also allowing for reduction of animals per group considering the 3R principles of animal welfare^45,46^. However, when applying longitudinal *in vivo* micro-CT, the effect of repeated anesthesia, handling and radiation associated with the scans on the study’s main outcome parameters and the general well-being of the animals have to be considered.

In this study, isoflurane was used as anesthetic agent for the *in vivo* micro-CT scans, which is the most commonly used inhalation anesthetic in longitudinal imaging studies with a fast on- and offset of anesthesia and low metabolism rate^17^. Despite its general use, isoflurane has been associated with several adverse effects, such as hepatic degenerative changes^47^, immunomodulation^48^, oxidative DNA damage^49^ as well as alterations in expression profiles of oncogene and tumor suppressor genes in the bone marrow^50^, which might potentially also affect fracture healing and the general well-being of the animals. A recent study by Hohlbaum et al. (2017)^17^, which assessed the impact of repeated isoflurane anesthesia (6 times for 45 min at an interval of 3–4 days) and the associated handling on the well-being of adult C57BL/6JRj mice, categorized the degree of distress as mild according to the EU Directive 2010/63 with only short-term impairment of well-being, mainly in the immediate post-anesthetic period. Therefore, isoflurane-based inhalation anesthesia and the associated handling of the animal is suggested for longitudinal studies with multiple anesthetic sessions per animal.

In this study, the applied micro-CT settings (55 kVp, 145 µA, 350 ms integration time, 500 projections per 180°, 21 mm field of view (FOV), scan duration ca. 15 min) were adapted from well-established protocols used for longitudinal monitoring of bone adaptation and implant integration in tail vertebrae in mice^14,51^. These settings are similar to protocols used in 2 of the 5 longitudinal fracture healing studies published so far^3,4^, while the 3 other studies did not specify the CT settings^2,32,33^. The radiation dosage (CT dose index, CTDI) associated with each scan in the current study was previously estimated to be 0.67 Gy per scan^51,52^. A radiation control experiment using similar settings did not show detrimental effects of 5 weekly *in vivo* micro-CT scans on bone development in tail vertebrae of mice^52^.

However, some studies in non-fractured bone indicate cumulative radiation-associated effects on bone properties particularly when using high radiation doses (>2G)^19^. The radiation effects also depend on animal age^19,27^ and ovariectomy (OVX)^11^, with younger and ovariectomized animals being more susceptible to radiation. So far, only one study assessed radiation-associated effects of longitudinal *in vivo* micro-CT on bone healing in a uni-cortical tibia burr-hole defect model, and did not find significant radiation effects on bone properties^1^. In this study, similar CT settings (45kVp, 133 µA, 200 ms integration time, 1000 projections per 180°) were used; however, due to the smaller dimensions of the defect and resulting shorter scanning time, the radiation dosage (0.36 Gy) was lower compared to our protocol (0.64 Gy).

In the current study, we assessed the combined imaging-associated impact (anesthesia, handling, radiation; 1x/week for 6 weeks) on callus formation and remodeling in a diaphyseal femur defect model in adult female C57BL/6J mice. We did not see any significant imaging-associated changes in bone volume and turnover in the fracture callus during the remodeling phase (post-operative week 5 and 6). Furthermore, no significant differences in callus mineralization between scanned (d0, week1-6) and control animals (d0, week 5+6) in any of the assessed callus volumes (TOT, DC, DP, FP) were observed. The distribution of the osseous callus volume into the three sub-volumes was also not significantly different between groups, indicating similar healing patterns. In addition, when separately assessing the cortical fragments between the inner pins (FC), also no significant differences in any of the assessed parameters were detected between groups. These 3D micro-CT based results were supported by 2D histomorphometric analysis, which also showed no significant differences in tissue composition (bone, cartilage, fibrous tissue, bone marrow) in the former defect region. In respect to clinical healing outcome, the percentage of unions in animals above the sub-critical defect length was 67% in both groups with restoration of the medullary cavity. Defects without cortical bridging only showed small cartilage residuals (<1%) as visualized by Safranin-O staining, indicating manifestation of non-union formation. In the defect regions of unions, no cartilage was seen, indicating progression towards the end of the healing period.

So far only few studies have focused on the underlying mechanisms of radiation-induced effects on bone mainly focusing on changes in expression profiles of osteoblast and osteoclast differentiation markers^30,35^. Recently, three consecutive *in vivo* studies by Chandra et al.^36,53,54^ showed, that radiation‐induced bone loss^53,54^ is mediated via Sclerostin inhibition of osteoanabolic Wnt-signaling^36^. First, they showed that focal radiation increases Sclerostin expression in trabecular bone in mice. Using a Sclerostin antibody, they were then able to block the radiation-induced deterioration in structural bone properties and consistently, trabecular bone in Sclerostin null mice was resistant to radiation. Based on these comprehensive analyses, we also assessed Sclerostin by immunohistochemistry, but did not see any differences in number of stained osteocytes, staining intensity and pattern between the scan and the control group in any of the assessed callus regions.

In summary, we did not see significant differences in a total of 10 assessed CT and histological parameters in the total volume (TOT) between the inner pins of the fixator as well as in the different callus sub-volumes (DC, DP, FP) and the adjacent cortical fragments (FC) between the scan and the control group. The sub-volume specific analysis also allowed capturing of distinct features only present in one region (e.g. significant post-operative increase in bone resorption in FC), that showed contrary progression compared to the overall picture (significant post-operative increase in bone formation in DC, DP and FP). This also shows the importance of VOI selection. The other longitudinal fracture healing studies either assessed a VOI consisting of the defect as well as adjacent bone fragments, or then only the callus in the defect region including the last slice of intact cortex^2,3^, or the callus in the defect and periosteal region combined^4^ was evaluated. An overall VOI including the defect region and cortical fragments can give an overview of callus characteristics, but sub-volume-specific differences might be missed. Therefore, analysis of different endosteal and periosteal callus regions as well as the adjacent cortical volume is favorable.

The current study has several limitations. We based our evaluation of longitudinal time-lapsed *in vivo* imaging for monitoring fracture healing on micro-CT and histological assessment of callus properties. Histomorphometry confirmed the micro-CT results. However, contouring of non-union defects is challenging and might be user dependent as indicated previously^55^. This could be addressed in future studies by implementing 2D-3D registration approaches^56^, allowing to transfer the automatically computed VOI borders generated for the micro-CT evaluation to the 2D histology analyses. We did not assess systemic effects associated with the repeated *in vivo* micro-CT measurements and isoflurane anesthesia. Due to the required post-operative scan defined in our ethical license, we did not include a second control group only receiving one micro-CT measurement at the endpoint. In respect to the underlying mechanisms associated with radiation effects on fracture healing, we chose to assess Sclerostin by immunohistochemistry, due to the comprehensive available literature. As proposed by Chandra et al.^36^, future studies could extend the analysis to other Wnt signaling inhibitors (e.g. DKK1) with the potential to find further targets for prevention of radiation-mediated bone loss. As we only included female animals in the study, our findings could be affected by the estrous cycle of the animals, which in future could be addressed by staging of the cycle using vaginal smears^57^. We minimized the risk of cycle-associated variance by using individual ventilated cages, which have been associated with the Lee–Boot effect (suppression or prolongation of estrous cycles) in group-housed adult female mice isolated from male animals^58^. In addition, recent literature indicates that unstaged female rodents are no more variable than males (revised in ^59^).

Despite these limitations, the longitudinal *in vivo* micro-CT based approach established in this study allows monitoring of the different healing phases in mouse femur defect models without significant anesthesia-, handling- and radiation-associated effects on callus properties. By registering consecutive scans of the defect region for each animal and implementing a two-threshold approach, data on bone turnover and mineralization kinetics can be obtained, which is important for targeting impaired callus remodeling under different conditions (e.g. premedication with bisphosphonates) with pharmacological and mechanical therapies. Importantly, repeated anesthesia, handling and radiation associated with the scans did not impair callus formation and remodeling. Therefore, this study supports the application of longitudinal *in vivo* micro-CT for healing-phase specific monitoring of fracture repair in mice. Further studies should evaluate the potential of this micro-CT-based monitoring approach for healing phase-specific discrimination of normal and impaired healing conditions.

## Methods

### Study design

A micro-CT based approach for longitudinal *in vivo* monitoring of fracture healing was established for a mouse femur defect model (for details on study design see Supplementary Table S1 online). All mice received a femur defect and post-operative micro-CT scans (vivaCT 40, ScancoMedical, Brüttisellen, Switzerland) were performed. The scan group then received weekly scans of the defect area (weeks 1-6). To assess the combined effect of radiation, anesthesia and handling associated with weekly micro-CT measurements, controls were only scanned after 5 weeks. Control animals then received another scan in post-operative week 6 to enable the assessment of both, static and dynamic bone parameters during the final remodeling phase of fracture healing.

### Animals

All animal procedures were approved by the authorities (licence number: 36/2014; Kantonales Veterinäramt Zürich, Zurich, Switzerland). We confirm that all methods were carried out in accordance with relevant guidelines and regulations (ARRIVE guidelines and Swiss Animal Welfare Act and Ordinance (TSchG, TSchV)). To study adult fracture healing female 12 week-old C57BL/6J mice were purchased from Janvier (Saint Berthevin Cedex, France) and housed in the animal facility of the ETH Phenomics Center (EPIC; 12h:12h light-dark cycle, maintenance feed (3437, KLIBA NAFAG, Kaiseraugst, Switzerland), 5 animals/cage) for 8 weeks. At an age of 20 weeks, all animals received a femur defect by performing an osteotomy (group 1: control group, defect length - 1.45mm ± 0.16, n=8; group 2: scan group, defect length −1.47mm ± 0.16, n=11; housing after surgery: 2-3 animals/cage; for details on study design see Supplementary Table 1 online). All defect surgeries were performed by the same veterinarian. Perioperative analgesia (25 mg/L, Tramal®, Gruenenthal GmbH, Aachen, Germany) was provided via the drinking water two days before surgery until the third post-operative day. For surgery and micro-CT scans, animals were anesthetized with isoflurane (induction/maintenance: 5%/1-2% isoflurane/oxygen). Perioperative handling and monitoring was performed by the surgeon and one other veterinarian.

### Femur osteotomy

In all animals an external fixator (Mouse ExFix, RISystem, Davos, Switzerland; stiffness: 24N/mm) was positioned at the craniolateral aspect of the right femur and attached using four mounting pins. First, the most distal pin was inserted approximately 2mm proximal to the growth plate, followed by placement of the most proximal and the inner pins. Subsequently, a femur defect was created using 2 Gigli wire saws.

### Time-lapsed *in vivo* micro-CT

Immediate post-surgery correct positioning of the fixator and the defect was visualized using a vivaCT 40 (Scanco Medical AG, Brüttisellen, Switzerland) (isotropic nominal resolution: 10.5 µm; 2 stacks of 211 slices; 55 kVp, 145 µA, 350 ms integration time, 500 projections per 180°, 21 mm field of view (FOV), scan duration ca. 15 min). Subsequently, the fracture callus and the adjacent bone between the inner pins of the fixator were scanned weekly using the same settings. Scans were registered consecutively using a branching scheme (registration of whole scan for bridged defects; separate registration of the two fragments for unbridged defects)^5^. Subsequently, morphometric indices (bone volume - BV, bone volume/total volume – BV/TV, bone formation rate – BFR, bone resorption rate - BRR) were computed (threshold: 395mg HA/cm^3^; for details on methods see ^5^). To assess mineralization progression, a second threshold (645mg HA/cm^3^) was applied and the ratio between highly and lowly mineralized tissue (BV_645_/BV_395_) was calculated. The two selected thresholds are included in our recently developed multidensity threshold approach^5^. Similar thresholds have previously been used in fracture healing studies^60,61^. The lower threshold (395mg HA/cm^3^) was selected to capture early callus formation, based on its previous application for assessment of lowly mineralized tissue in the metabolically active trabecular compartment^62,63^. The choice of the second threshold (645 mg HA/cm^3^) for monitoring callus mineralization was based on thresholds previously applied to well-mineralized cortical bone^62,63^. It has to be considered, that micro-CT-based assessment of the mineralization process is known to be affected by the polychromatic cone beam (≤10% error in the quantification of the local bone mineral density^64^). We used an aluminium filter to further reduce the effect of beam hardening^65^ and applied scanning settings previously evaluated for monitoring *in vivo* mineralization kinetics^66^. According to the standard clinical evaluation of X-rays, the number of bridged cortices per callus was evaluated in two perpendicular planes (UCT Evaluation V6.5-1, Scanco Medical AG, Brüttisellen, Switzerland). A ‘‘healed fracture’’ was considered as having a minimum of at least three bridged cortices per callus. For evaluation, four volumes of interest (VOIs) were defined, which were created automatically from the post-operative measurement (Fig. 1): defect center (DC), defect periphery (DP), cortical fragment center (FC), and fragment periphery (FP). Data were normalised to the central VOIs: DC/DC, DP/DC, FC/FC, FP/FC.

### Histology

Histological analyses were performed in a sub-set of animals (n=6/group). On day 42, femora were excised, the femoral head was removed and the samples were placed in 4% neutrally buffered formalin for 24 hours and subsequently decalcified in 12.5% EDTA for 10-14 days. The samples were embedded in paraffin and 4.5 µm longitudinal sections were stained with Safranin-O/Fast Green: Weigert’s iron haematoxylin solution (HT1079, Sigma-Aldrich, St. Louis, MO) - 4min, 1:10 HCl-acidified 70% ethanol - 10s, tap water - 5min, 0.02% Fast Green (F7258, Sigma-Aldrich, St. Louis, MO) - 3min, 1% acetic acid - 10s, 0.1% Safranin-O (84120, Fluka, St. Louis, MO) - 5min. Images were taken with Slide Scanner Pannoramic 250 (3D Histech, Budapest, Hungary) at 20x magnification.

For immunohistochemical staining of Sclerostin, nonspecific sites were blocked (1% BSA/PBS + 1% rabbit serum) for 60 min at room temperature. Subsequently, the sections were incubated with the primary antibody against Sclerostin (AF1589, R&D Systems, Minneapolis, MN; 1:150 in 1%BSA/PBS + 0.2% rabbit serum) overnight at 4°C. To detect the primary antibody, a secondary biotinylated rabbit anti-goat-IgG antibody (BAF017, R&D Systems, Minneapolis, MN) was added for 1 h at room temperature. For signal amplification, the slides were incubated with avidin-biotin complex (PK-6100 Vector Laboratories, Burlingame, CA) for 30 min. Diaminobenzidine (Metal Enhanced DAB Substrate Kit, 34065 ThermoFisher Scientific, Waltham, MA) was used as detection substrate. Counterstaining was performed with FastGreen (F7258, Sigma-Aldrich, St. Louis, MO). Species-specific IgG was used as isotype control. Images were taken with Slide Scanner Pannoramic 250 (3D Histech, Budapest, Hungary) at 40x magnification.

For histomorphometric analysis, the callus and adjacent cortical fragments between the inner pins of the fixator were manually contoured on Safranin-O stained sections and the tissue area of this region of interest (ROI) was quantified using commercial software (n=6/group; Fiji^67^). Similarly, the different tissue types (bone, cartilage, fibrous tissue and bone marrow) in the ROI were manually contoured and quantified.

### Statistics

CT analysis and histomorphometry: Data were tested for normal distribution (Shapiro-Wilk-Test) and homogeneity of variance (Levene-Test). Depending on the test outcome, group comparisons (scan versus control group) of data derived at single time points were performed by two-tailed Student’s t-test or Mann-Whitney U-test (IBM SPSS Statistics Version 23). For statistical evaluation of repeated measurements (scan group) dependent on results from normality and variance tests, either one-way analysis of variance (ANOVA) with Bonferroni correction or Friedman test with Dunn correction for multiple comparisons (GraphPad Prism 7) were performed. Two-way ANOVA was used for longitudinal comparison of the body weight between the control and the scan group. The level of significance was set at p < 0.05.

## Acknowledgements

The authors gratefully acknowledge support from the EU (BIODESIGN FP7-NMP-2012-262948 and ERC Advanced MechAGE ERC-2016-ADG-741883). E. Wehrle received funding from the ETH Postdoctoral Fellowship Program (MSCA-COFUND, FEL-25_15-1).

## Author Contributions Statement

The study was designed by E.W., G.A.K., S.H. and R.M.. The experiments were performed by E.W., G.A.K. and A.C.S.. Data analyses were performed by E.W. and D.C.B.. The manuscript was written by E.W. and reviewed and approved by all authors.

## Data availability

All necessary data generated or analyzed during the present study are included in this published article and its Supplementary Information files (preprint available on BioRxiv (BIORXIV/2019/692343). Additional information related to this paper may be requested from the authors.

