## Supplementary figures and images for "Evaluation of longitudinal time-lapsed *in vivo* micro-CT for monitoring fracture healing in mouse femur defect models"

### Supplementary Figure S1

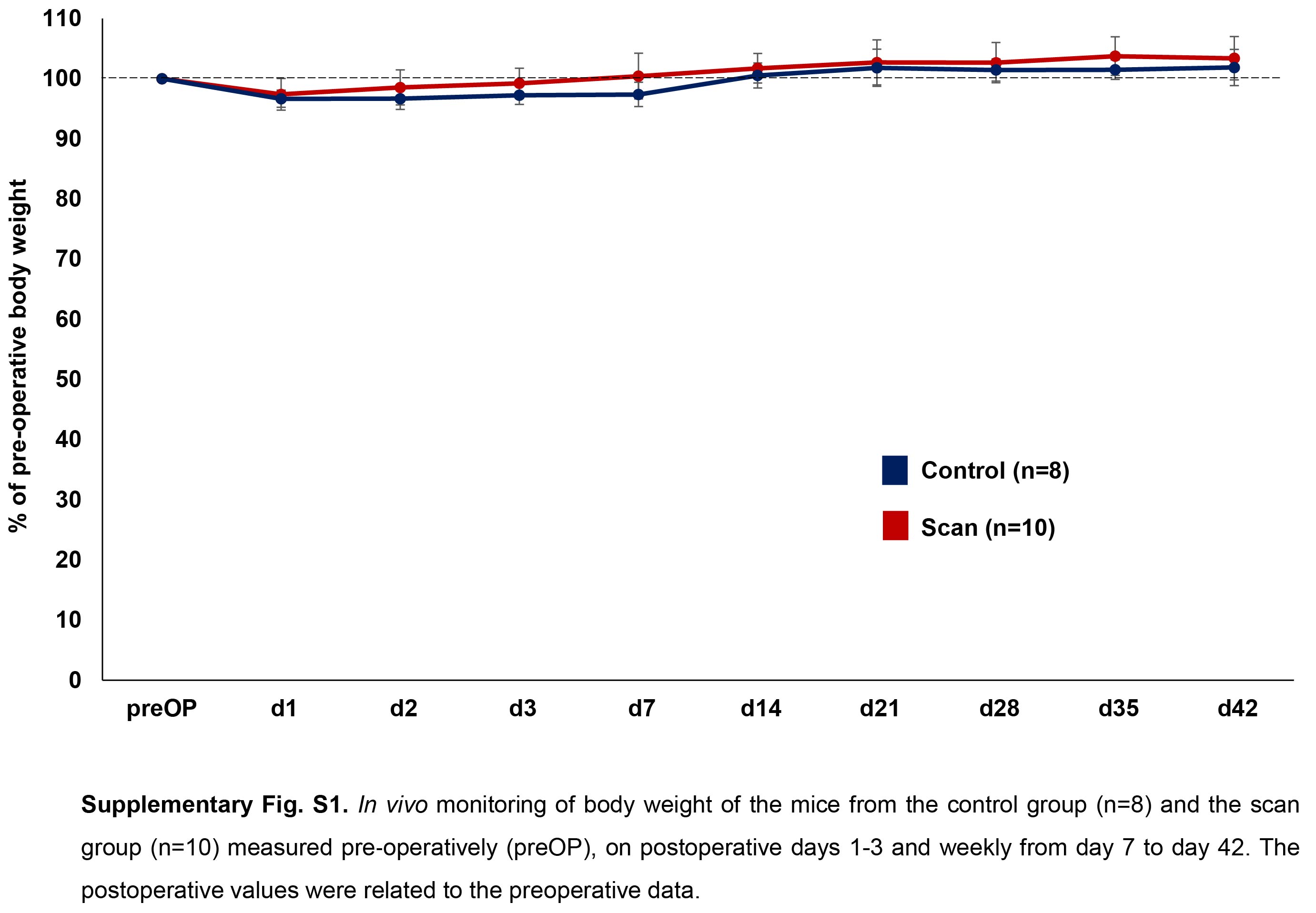

### Supplementary Figure S2

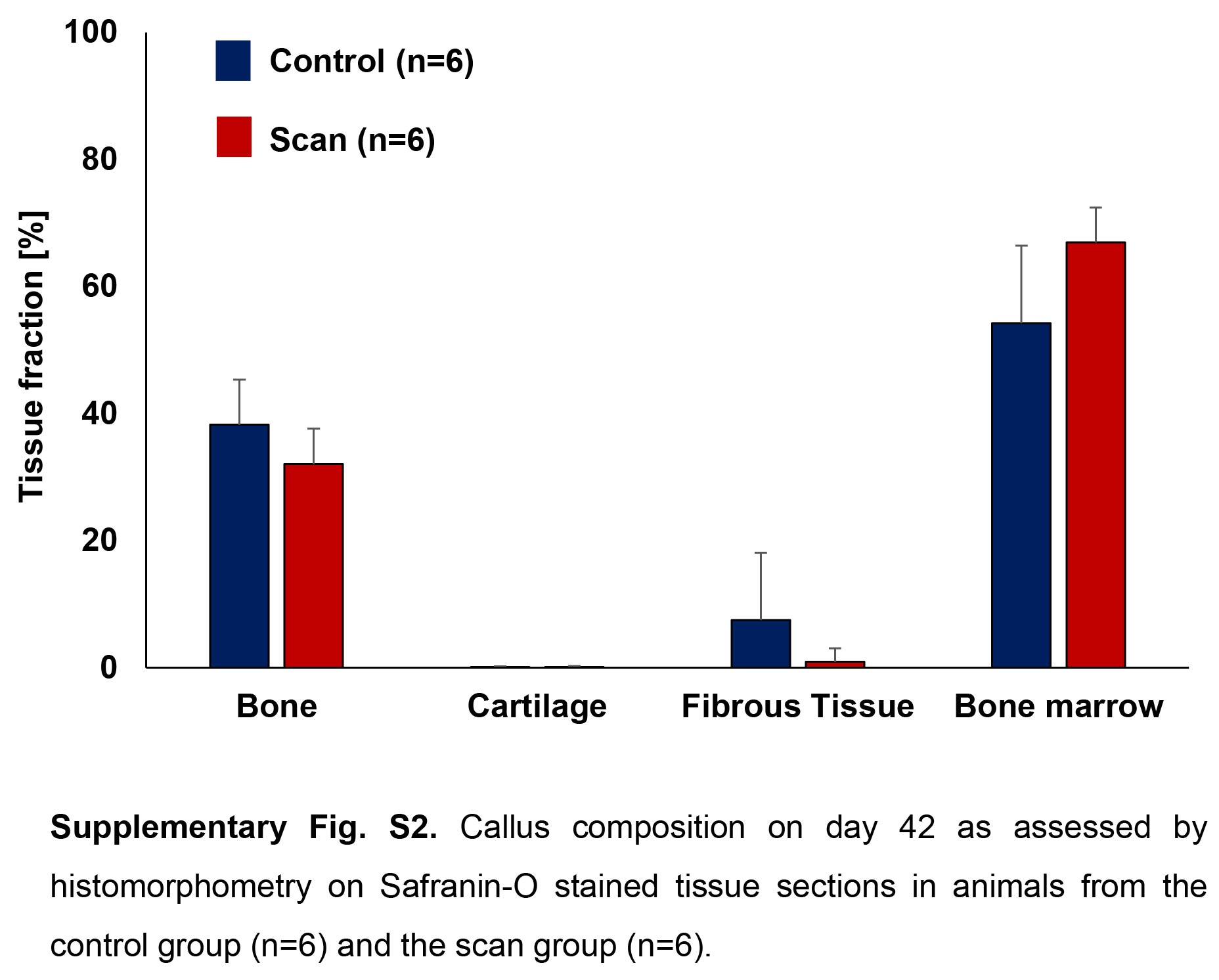
