## Supplementary Table S1 for "Evaluation of longitudinal time-lapsed *in vivo* micro-CT for monitoring fracture healing in mouse femur defect models"

**Supplementary Table S1.** Study design (female 20 week-old C57BL/6J mice)

| Group | Group size | Femur defect (at 20 weeks of age) | *In vivo* micro-CT measurements | Registration of micro-CT scans # | Histology |
|---|---|---|---|---|---|
| Scan | n=11 | x (n=11) | d0, week 1-6 (n=11) | week 1-6 to week 0-5 (n=10) | week 6 (n=6) |
| Control | n=8 | x (n=8) | d0, week 5+6 (n=8) | week 6 to week 5 (n=8) | week 6 (n=6) |

### micro-CT scan taken at timepoint x registered to micro-CT scan taken at timepoint x-1
